# The adaptive potential of non-heritable somatic mutations

**DOI:** 10.1101/2021.04.30.442123

**Authors:** Paco Majic, E. Yagmur Erten, Joshua L. Payne

## Abstract

Non-heritable somatic mutations are typically associated with deleterious effects such as in cancer and senescence, so their role in adaptive evolution has received little attention. However, most somatic mutations are harmless and some even confer a fitness advantage to the organism carrying them. We hypothesized that heritable, germline genotypes that are likely to express an advantageous phenotype via non-heritable somatic mutation will have a selective advantage over other germline genotypes, and this advantage will channel evolving populations toward more fit germline genotypes, thus promoting adaptation. We tested this hypothesis by simulating evolving populations of developing organisms with an impermeable germline-soma separation navigating a minimal fitness landscape. The simulations revealed the conditions under which non-heritable somatic mutations promote adaptation. Specifically, this can occur when the somatic mutation supply is high, when only very few cells with the advantageous somatic mutation are required to increase organismal fitness, and when the somatic mutation also confers a selective advantage to cells with that mutation. We therefore provide proof-of-principle that non-heritable somatic mutations can promote adaptive evolution via a process we call somatic genotypic exploration. We discuss the biological plausibility of this phenomenon, as well as its evolutionary implications.

**Significance:** The immensity of non-heritable genetic diversity arising in the soma has been largely disregarded in evolutionary theory as a source of adaptation. Here, we introduce a model in which non-heritable somatic mutations arising during development confer an organismal fitness advantage. Analysis of this model shows how such mutations channel evolving populations toward adaptive germline genotypes. This is most likely to occur when somatic mutations confer a fitness benefit at both the cellular and organismal levels, evoking a synergistic form of multi-level selection that contrasts with the antagonistic forms typically associated with somatic mutations in cancer and senescence. As such, our study invites a new view of somatic genetic diversity in evolutionary theory as a potential source of adaptation.

## Introduction

During the development of most animals, an early distinction occurs between the germline – the population of cells that are fated to differentiate into gametes – and the soma – the cells composing the rest of the body. August Weismann noted that any variation arising in the soma during the lifetime of an organism would be temporary and non-heritable, because it would not be present in the reproductive cells [1]. The non-heritability of somatic variation weakened Lamarckian arguments concerning the role of acquired variation in adaptation and set the stage for a neo-Darwinian take on evolution [2, 3]. Within this paradigm, the somatic organism came to be viewed as a mere “excrescence” [4] or a “dead-end replicator” [5], and the non-heritable genetic variation arising in it as an evolutionary cul-de-sac [5–8]. Consequently, studies of somatic mutation in animal evolution mainly focused on checks against their deleterious effects at the organismal level, such as in cancer and senescence [9–14].

A strong research focus on the molecular biology of cancer has unveiled aspects of somatic mutations that go beyond their deleterious effects. Somatic mutations are ubiquitous and, more often than not, harmless [15–18]. They are detected at different frequencies within the soma, partly depending on whether they arise early or late during development [19–22], and partly because of the selective competitiveness of mutant cells [17, 23–25], since somatic mutations can increase in frequency within the body when they confer a higher proliferative potential or lower mortality to the cells carrying them [26]. Although the clonal expansion resulting from this somatic selective process is one of the characteristics of cancer, positive selection of cells with somatic mutations can also occur without causing any apparent disease phenotypes in the tissues containing the mutant cells [15, 17, 24, 25, 27, 28].

In some cases, somatic mutations are beneficial not only for individual cells, but also for the organism with the mutated cells. A classic example is the adaptive immune system of jawed vertebrates, in which somatic mutants are selected within the body based on their affinity to the pathogens they help deter [29, 30]. Somatic mutations can be beneficial in other physiological contexts as well, such as in the liver, where they can promote regeneration after injury, protect against toxins, and prevent malignant transformation [31]. They can also ameliorate the consequences of deleterious mutations that cause hematopoietic diseases [32] or serious developmental syndromes [33]. In such cases, somatic mutations may facilitate the persistence of deleterious germline mutations by buffering their negative effects on organismal fitness until a compensatory or reversion mutation arises in the germline [16].

Research on the evolutionary consequences of beneficial somatic variation has mainly focused on plants [34–38]. However, plants and other organisms for which an adaptive potential of somatic mutation has been considered (e.g., corals [39] and red algae [40]) do not always produce somatic variation in the strict sense of Weismann – either because there is a blurry germline-soma distinction or because they can reproduce clonally, which often makes somatic variation heritable [38, 41–43]. There is, however, evolutionary potential in strictly non-heritable somatic mutation. Under a scenario in which non-heritable somatic mutations can confer a fitness advantage to the organism carrying them, selection can act on the potential to acquire such mutations, as evidenced by the evolution of mechanisms that direct or intensify the production of somatic genetic variation [44–48]. These mechanisms tend to target specific genomic regions and function in specialized cell types, and what ultimately gets selected is the mechanisms producing the somatic mosaicism rather than the somatic mutant genotypes themselves [8, 34, 44, 46, 49].

We envision a complementary, general model in which selection acting on genotypes with a potential to acquire non-heritable somatic mutations that are beneficial to the organism facilitate adaptation, even in the absence of a mechanism to intensify somatic diversity. Given the sheer number of cells in the soma and their increased mutation rates relative to the germline [50–53], we reason that beneficial mutations often first arise in the soma. Similar to so-called phenotypic mutations [54], which arise due to errors in transcription or translation, heritable genotypes that are similar in sequence to a non-heritable beneficial genotype may occasionally confer the fitness benefit of the non-heritable beneficial genotype to an organism via somatic mutation. Placing this model in the context of an adaptive landscape [55], the germline genotype can be one or more mutations away from an adaptive peak, and somatic mutations can confer a fitness benefit to an organism by producing non-heritable beneficial genotypes that are near or atop the adaptive peak. This can cause a smoothing of the landscape [56], which may promote adaptation towards an adaptive peak by increasing the probability that the beneficial mutation arises in the germline, because genotypes that are near the adaptive peak are more likely to be selected. We thus hypothesize that non-heritable somatic mutations expressing a fitness advantage may channel evolving populations towards adaptive peaks, thus promoting adaptation.

## Results

### Model overview

To study the potential of non-heritable somatic mutations to promote adaptation, we modelled an evolving population of *N* multicellular organisms with an impermeable germline-soma separation navigating a minimal fitness landscape. We used a two-locus, two-allele model with alleles a and A for the first locus and b and B for the second locus, to represent a landscape with a single adaptive peak at genotype AB, which confers a selective advantage *s*_organism_ to the organism relative to the other genotypes in the landscape (Fig. 1A, Methods). We simulated evolution with non-overlapping generations, such that the initial population was composed exclusively of organisms with the ab genotype. Each generation consisted of a developmental phase followed by a reproductive phase (Fig. 1B, Methods). In the developmental phase, the soma of each individual developed from a single cell with a given zygotic genotype in *D* developmental cycles until reaching the final somatic size of 2^*D*^ cells. For each cell division, somatic mutations occurred at rate *μ*_soma_ per locus. The genotypic composition of the soma at the end of development defined organismal fitness. As such, we allowed somatic mutations only during development and did not model the mature lifespan of organisms. In the reproductive phase, organismal reproductive success was proportional to fitness, and germline mutations occurred at rate *μ*_germline_ per locus.

**Figure 1.**
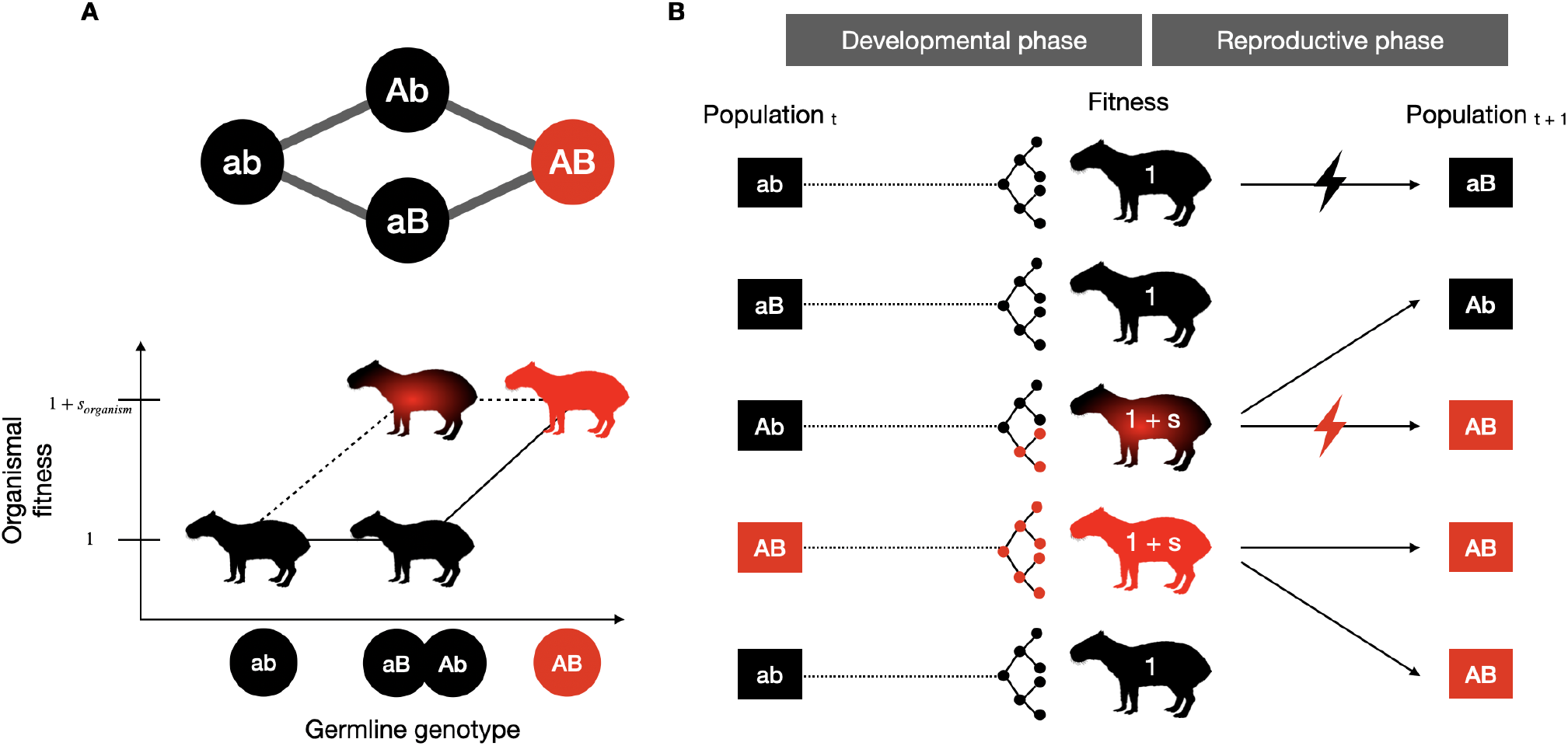
Baseline model. A) Fitness landscape represented as a two-locus two-allele model. Each line connecting genotypes in black and red nodes corresponds to a single mutational step. Genotype AB (red; peak genotype) confers a higher fitness to its carrier, whereas all other genotypes confer no selective advantage. The Cartesian axes represent the individual fitness value as a function of the distance of the heritable germline genotype to the peak. Solid lines indicate the case in which somatic mutations are either not present or produce no selective advantage, such as in our control simulations. Dashed lines show how beneficial somatic mutations can increase organismal fitness. B) Representation of a single generation in our simulations. At generation *t*, individuals in a population of size *N* enter a developmental phase. During this phase, starting from a single cell with each individual’s germline genotype, *D* developmental cycles occur until the final somatic size 2^*D*^ is reached. At each developmental cycle, somatic mutations occurring at rate *μ*_soma_ can modify the distance to the peak of each somatic cell. Based on a fitness function, at the end of the developmental phase, the final genotypic composition of the soma defines the fitness of each individual. During the reproductive phase, the population is sampled based on the individual fitness values to create a new population for the next generation. Before entering the developmental phase of generation *t* + 1, germline mutations may occur at rate *μ*_germline_, represented by red and black lightning bolts in our diagram.

### Non-heritable somatic mutations can promote adaptation

We ran two versions of our model across a range of somatic and germline mutation rates. In the first version, somatic mutations did not confer an organismal selective advantage. This served as a control and as a point of comparison with traditional population genetic models that disregard somatic development. We implemented this version of our model by simulating each generation using only the reproductive phase, thus ignoring somatic mutations that could arise during the developmental phase. Fitness was therefore defined by the germline genotype alone. In the second version, somatic mutations conferred an organismal selective advantage. We implemented this version of our model using a fitness function in which an individual attained the full selective advantage *s*_organism_ if at least one somatic cell had the peak genotype at the end of the developmental phase – an assumption we later relax. Fitness was therefore defined by the somatic composition of the organism.

Fig. 2A-D shows the evolutionary outcomes of these simulations for a range of germline and somatic mutation rates. In the control simulations, the mean fitness of the population increased with the germline mutation rate, after it exceeded a threshold of approximately 5×10^−7^ mutations per locus per generation (Fig. 2A). Under these high germline mutation rates, populations increased in fitness by converging on the peak genotype via stochastic tunneling (Fig. 2C) [57]. When somatic mutations conferred a selective advantage to the organism, the mean fitness of the population increased with both the germline and somatic mutation rates, thus expanding the parameter space in which higher fitness evolved, relative to the control simulations (compare Figs. 2A and 2B). For low somatic mutation rates, this expansion was explained by a decreased threshold on the germline mutation rate past which populations converged on the peak genotype (Fig. 2D). In contrast, for higher somatic mutation rates, fitness increases were not due to convergence on the peak genotype in the germline. Rather, populations remained one or two mutations away from the adaptive peak (Fig. 2D), because the high somatic mutation rates all but guaranteed the emergence of the peak genotype in the soma, thus rendering non-peak germline genotypes selectively equivalent to peak germline genotypes. Additionally, beyond expanding the parameter space, somatic mutations accelerated the rate of evolution to the peak for germline mutation rates where populations converged on the peak germline genotype in both versions of our model (Fig. 2E).

**Figure 2.**
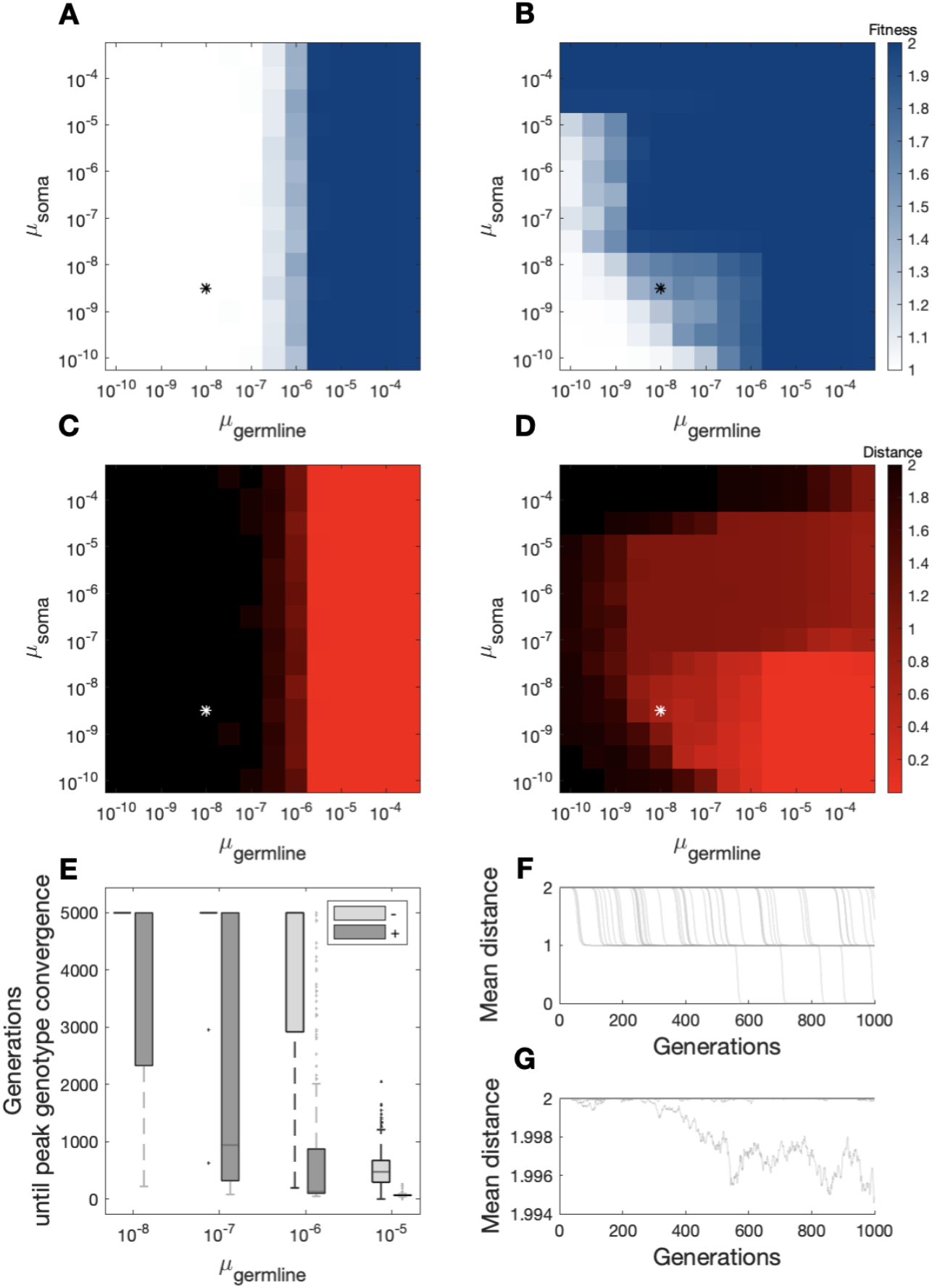
Non-heritable somatic mutations can promote adaptation. (A,B) Mean fitness of populations and (C,D) mean mutational distance to the peak after 5000 generations when somatic peak genotypes (A,C) did not provide an organismal selective advantage and (B,D) when they did, for a range of somatic mutation rates (mutations per locus per cell division) and germline mutation rates (mutations per locus per generation). The values shown are the mean across 500 replicates for each parameter combination. Asterisks indicate the combination of *μ*_soma_=5×10^−9^ and *μ*_germline_=1×10^−8^, which approximates empirically estimated mutation rates from mouse and human cells [52]. (E) Distribution of the number of generations required to converge on the peak genotype in 500 simulations for four different values of the germline mutation rate, when somatic peak genotypes provided a selective advantage (+) and when they did not (-). For all simulations in this panel, *μ*_soma_=5×10^−9^ mutations per locus per division. (F,G) Evolutionary trajectories over the first 1000 generations for 100 randomly chosen replicates when somatic peak genotypes (F) provided a selective advantage and (G) when they did not, under the mutation rates indicated by the asterisks in A-D. We ran all simulations with a population size *N* = 100000, a selective advantage *s*_organism_ = 1 and a number of developmental cycles *D* = 25.

We note that empirical mutation rates from mouse and human cells (*μ*_soma_ = 5×10^−9^, *μ*_germline_ = 1×10^−8^; [52]) fall within the range of mutations rates where somatic mutations can promote adaptation via convergence on the peak germline genotype (Fig. 2A-D, asterisk). For these mutation rates, an initial stage of drift was often followed by two consecutive selective sweeps (Fig. 2F). In contrast, in control simulations, genetic drift dominated the evolutionary dynamics, such that populations remained two mutations from the peak (Fig. 2G).

Taken together, these results provide proof-of-principle that non-heritable somatic mutations can promote adaptation, even under biologically realistic germline and somatic mutation rates.

### Somatic mutation supply determines evolutionary outcomes

The results presented above revealed three distinct evolutionary outcomes when somatic mutations improved organismal fitness, depending upon the somatic and germline mutation rates (Fig. 3A): (*i*) populations did not increase in fitness, and remained two mutations away from the peak; (*ii*) populations increased in fitness by converging on the peak; and (*iii*) populations increased in fitness, but remained one or two mutations away from the peak.

**Figure 3.**
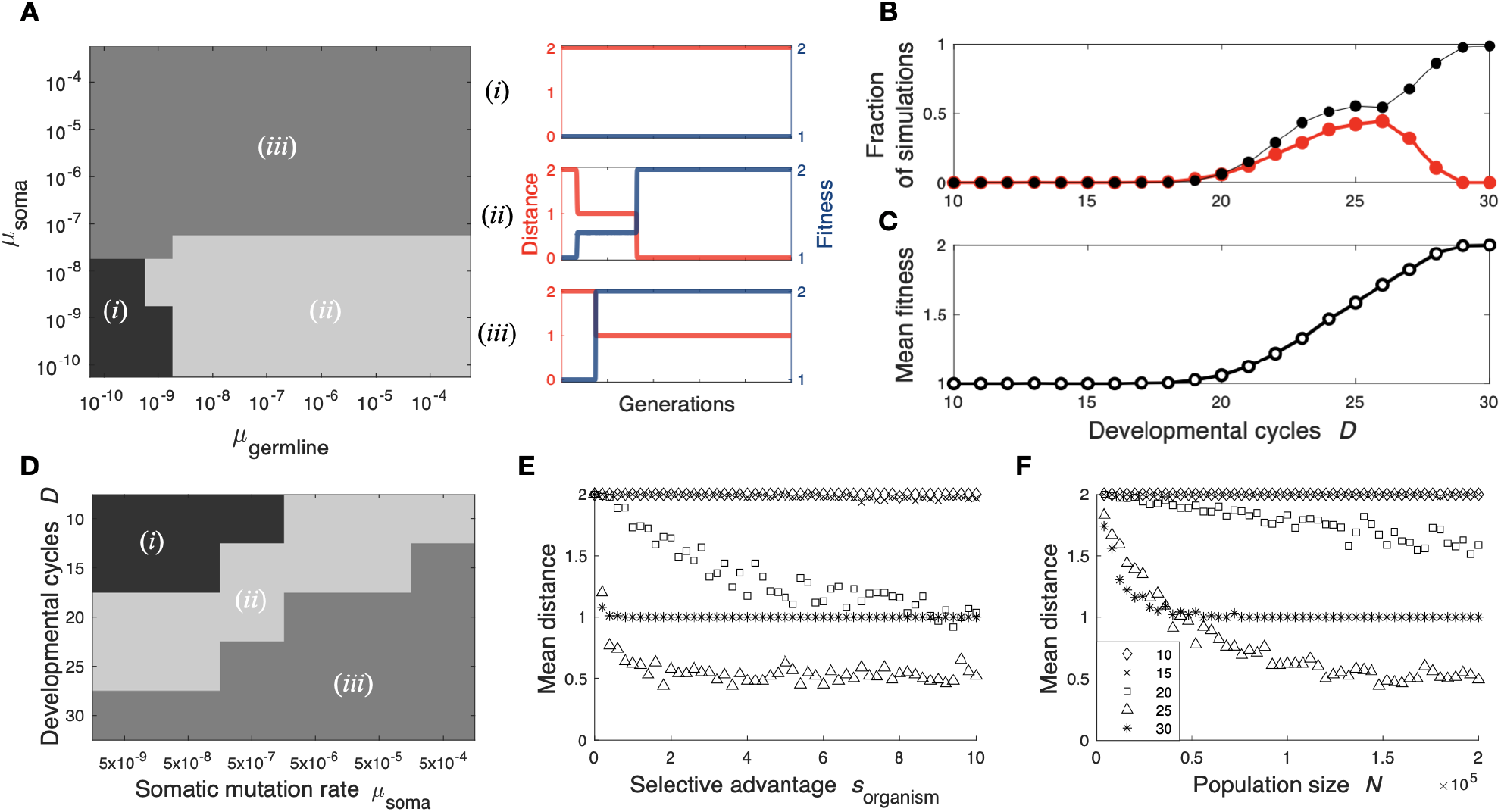
Evolutionary outcomes in relation to model parameters. (A) Three distinct evolutionary outcomes emerge in our model: (*i*) populations do not evolve the maximum fitness of 1 + *s*_organism_ after 5000 generations, (*ii*) populations evolve the maximum fitness of 1 + *s*_organism_ after 5000 generations and converge on the peak genotype, and (*iii*) populations evolve the maximum fitness of 1 + *s*_organism_ after 5000 generations, but do not converge on the peak genotype. Regions shaded according to the outcome realized by the simulations for each parameter combination. Subpanels to the right show the trajectories for mean distance to the peak (red) and mean fitness (blue) in representative simulations, for each of the three evolutionary outcomes. (B) The fraction of simulations in which the population reached the peak genotype (red) or remained one mutation away from the peak genotype (black) after 5000 generations is shown in relation to *D*. (C) The mean population fitness after 5000 generations of the same simulations as in (B). (D) Evolutionary outcomes *i*, *ii* or *iii* for different combinations of somatic mutation rates and developmental cycles. (E,F) Mean distance of the germline genotype to the peak after 5000 generations across a range of values for (E) selective advantage and (F) population size. The different symbols correspond to different number of developmental cycles *D*, as shown in the legend in panel (F). For (D-F), we performed 100 simulations for each combination of parameters. The baseline parameters were *N* = 100000, *s*_organism_ = 1, *μ*_soma_ = 5×10^−9^ and *μ*_germline_ = 1×10^−8^.

These results suggest that somatic mutation supply determines which evolutionary outcome emerges. Another factor that influences somatic mutation supply beside somatic mutation rate is the number of cells in the soma, which in our model is given by the number of developmental cycles *D*. To assess how somatic mutations influence adaptation for organisms of different size, we modified our baseline model to include a range from *D* = 10 to *D* = 30, which produces final somatic cell counts between 2^10^ = 1024 and 2^30^ = 1.07×10^9^, respectively — values that approximate the number of cells in tissues, organs, and entire animals (Table 1). Modifying the mutation supply in this way resulted in similar evolutionary outcomes as when varying somatic mutation rates (Fig. 3B,C). Specifically, after 5000 generations, no populations increased in fitness when *D* < 19, populations tended to increase in fitness by evolving the peak genotype when 19 ≤ *D* ≤ 28, and populations increased to a maximum fitness of 1 + *s*_organism_ without reaching the peak when *D* > 28. By varying somatic mutation rate together with the number of developmental cycles, we observed that the evolutionary outcome of our model depended on the interaction between these two factors of somatic mutation supply (Fig. 3D). For example, high somatic mutation rates facilitated convergence on the peak genotype even for populations of organisms with the smallest soma considered (*D* = 10, which approximates the somatic size of an adult *Caenorhabditis elegans*, Table 1).

**Table 1.**
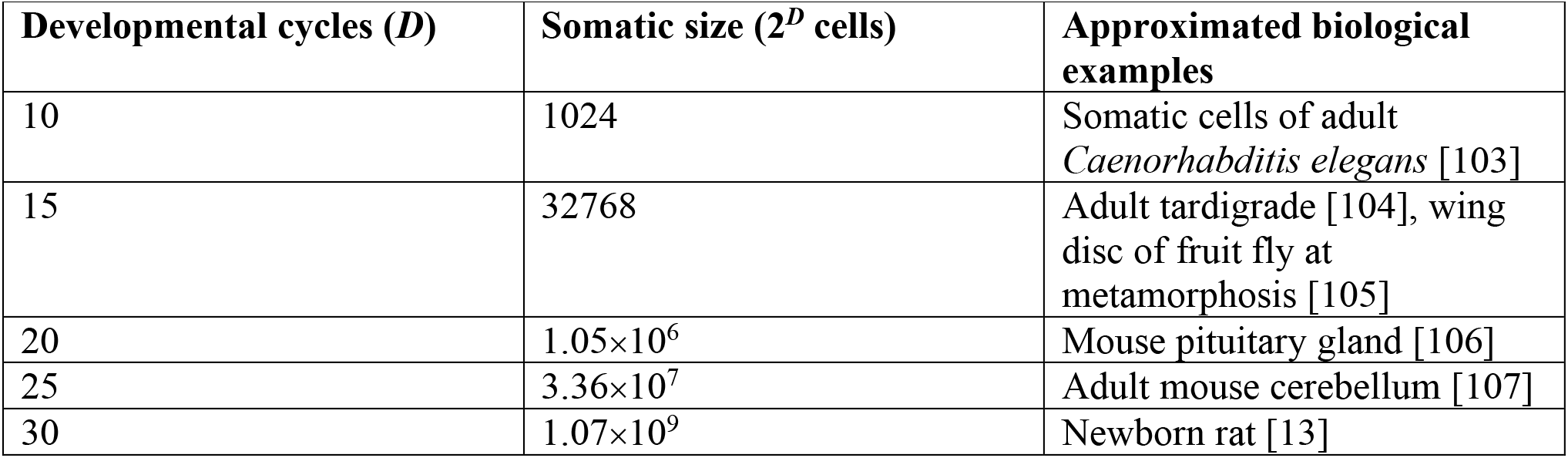
Number of cells across different biological examples

| Developmental cycles ($D$) | Somatic size ($2^D$ cells) | Approximated biological examples |
| --- | --- | --- |
| 10 | 1024 | Somatic cells of adult *Caenorhabditis elegans* [103] |
| 15 | 32768 | Adult tardigrade [104], wing disc of fruit fly at metamorphosis [105] |
| 20 | $1.05 \times 10^6$ | Mouse pituitary gland [106] |
| 25 | $3.36 \times 10^7$ | Adult mouse cerebellum [107] |
| 30 | $1.07 \times 10^9$ | Newborn rat [13] |

To explore the importance of somatic mutation supply relative to other factors affecting evolutionary dynamics in our model, we additionally varied the population size *N* and the selection coefficient *s*_organism_ of the AB genotype (Methods). The number of populations converging on the peak genotype increased as either of these parameters increased, but only for some intermediately sized somas (Fig. 3E,F). Thus, although population size *N* and selection coefficient *s*_organism_ can influence the probability with which a population converges on the peak germline genotype, it is ultimately the somatic mutation supply, defined by the somatic mutation rate and the size of the organism, that determines which of the three evolutionary outcomes emerge.

### Diminishing returns fitness functions restrict the adaptive potential of somatic mutations

So far, we have used a fitness function in which a single somatic cell with the peak genotype is sufficient to confer the full selective advantage *s*_organism_ to the organism. Relaxing this assumption to account for more realistic biological scenarios, we studied six diminishing returns fitness functions, in which organismal fitness depended on the fraction of cells with the peak genotype in an individual’s soma at the end of development (Fig. 4A; Methods). Figures 4B-D show the probability of converging on the adaptive peak in relation to the selection coefficient *s*_organism_ for three different somatic mutation rates using the six fitness functions. Under our baseline somatic mutation rate, beneficial somatic mutations only promoted adaptation for two of the fitness functions, specifically those that required the fewest somatic cells with the peak genotype to confer the full selective advantage (Fig. 4B). With these fitness functions, in a soma of 2^25^ cells, ~30 and ~300 cells with the peak genotype are needed to confer 10% of the selection coefficient *s*_organism_, respectively, and at least ~3000 and ~30000 cells (representing less than 0.01% of the total somatic cells) with the peak genotype are needed to confer the full selection coefficient *s*_organism_, respectively. For these fitness functions, increasing the somatic mutation rate two- and ten-fold increased the probability of converging on the adaptive peak for all values of *s*_organism_ (Fig. 4C,D), and for the ten-fold increase, non-heritable somatic mutations occasionally promoted adaptation for an additional fitness function (Fig. 4D). Thus, even under high somatic mutation rates and with high selection coefficients, somatic mutations are unlikely to promote adaptation if more than very few somatic cells with the peak genotype are required to confer the full selective advantage *s*_organism_.

**Figure 4.**
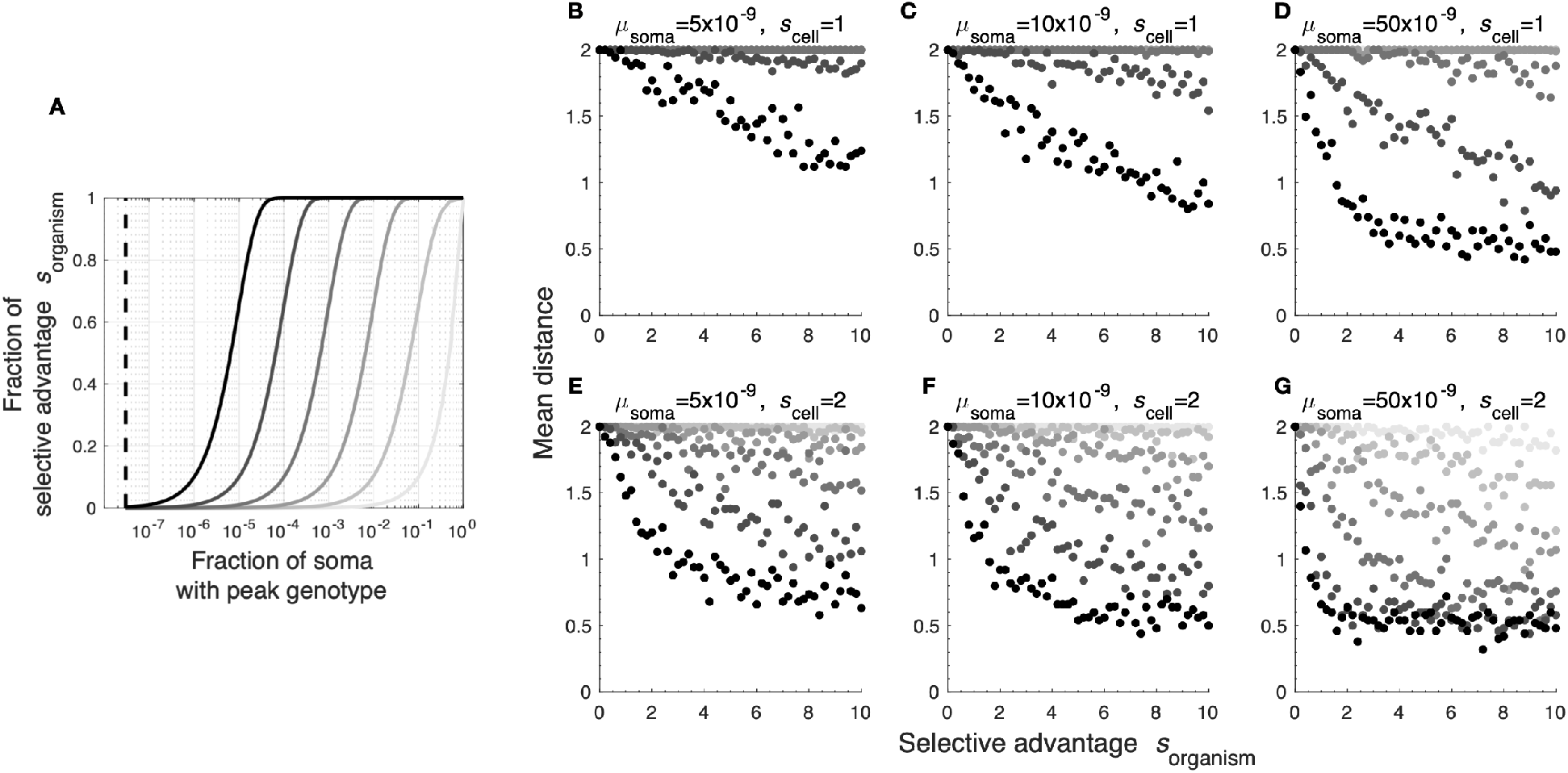
Influence of diminishing returns fitness functions and cell-lineage selection on the adaptive potential of somatic mutations. (A) Fraction of the selective advantage conferred to an individual as a function of the fraction of cells in the soma with the peak genotype. The six fitness functions are shown in different shades of gray; lighter gray indicates more restrictive fitness functions. The black dashed line indicates the value of 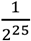, which is the minimum fraction of somatic cells with peak genotypes that are needed to confer the full selective advantage in the baseline model with *D* = 25 developmental cycles. (B-G) Mean distance to the peak in populations after 5000 generations across a range of selective advantages under each of the six different fitness functions from (A). Each point represents the mean across 50 replicate simulations for each parameter combination. For (E-G), we modified the simulations such that cells with peak genotypes had a fitness advantage *s*_cell_=2 over cells with other genotypes. We performed the simulations in (B-G) with the somatic mutation rates indicated above the panels, while the remaining parameters were *N* = 100000, *D* = 25 and *μ*_germline_ = 1×10^−8^.

### The adaptive potential of non-heritable somatic mutations under multi-level selection

We have so far assumed that somatic mutations confer a selective advantage to the organism, but not to the cell. Yet, cell-lineage selection is common in development and biases mosaic and chimeric cellular compositions [7, 37, 42, 58–63], helping to explain why some somatic mutations are recurrently detected across different individuals [64]. Cells can attain increased fitness if they better respond to signals in their developmental environment, make better use of resources, induce apoptosis of neighboring cells, proliferate more, or die less [65, 66]. We therefore included cell-lineage selection in our model, reasoning that it might expand the set of fitness functions under which non-heritable somatic mutations promote adaptation by increasing the number of cells with the peak genotype in the soma. We assumed somatic cells with peak genotypes had a cellular fitness advantage *s*_cell_ (Methods), but kept the final size of the soma at 2^*D*^ cells, inspired by systems with determinate growth [67]. In other words, we assumed that cells with a fitness advantage increased in frequency without affecting the final size of the organism. After applying these modifications to our model, we ran simulations using the same six fitness functions described above (Fig. 4A).

Doubling the cellular proliferative advantage of somatic peak genotypes (*s*_cell_ = 2) expanded the conditions under which non-heritable somatic mutations promoted adaptation, even for the lowest somatic mutation rate considered (Fig. 4E). Increasing the somatic mutation rate two- or ten-fold expanded the conditions even further (Fig. 4F,G), to the point that, even under the most restrictive fitness function considered (lightest shade of gray in Fig. 4A), some populations converged on the peak germline genotype (Fig. 4G). Thus, when a somatic mutation simultaneously benefits the organism and the somatic mutant cell within the context of development, non-heritable somatic mutations can promote adaptation across a broader range of fitness functions.

### Somatic mutations can facilitate fitness valley crossing

Thus far, we studied a fitness landscape with three genotypes of equal fitness (ab, Ab, and aB) and a fourth genotype (AB) with a selective advantage *s*_organism_ (Fig. 1A). We now study a rugged fitness landscape with two adaptive peaks separated by an adaptive valley. Specifically, we assigned a fitness of 1 to genotype ab, a fitness of 1 + *s*_1_ to genotype AB, and a basal fitness of 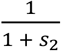 to the intermediate genotypes aB and Ab, which could be increased via somatic mutation to AB (Fig. 5A). Such landscape ruggedness can hinder evolution, because populations can become trapped in local adaptive peaks [68]. However, we reasoned that non-heritable somatic mutations might permit valley crossing, because the fitness disadvantage of the Ab or aB germline genotypes could be offset by the fitness advantage of somatic mutations to the AB genotype.

**Figure 5.**
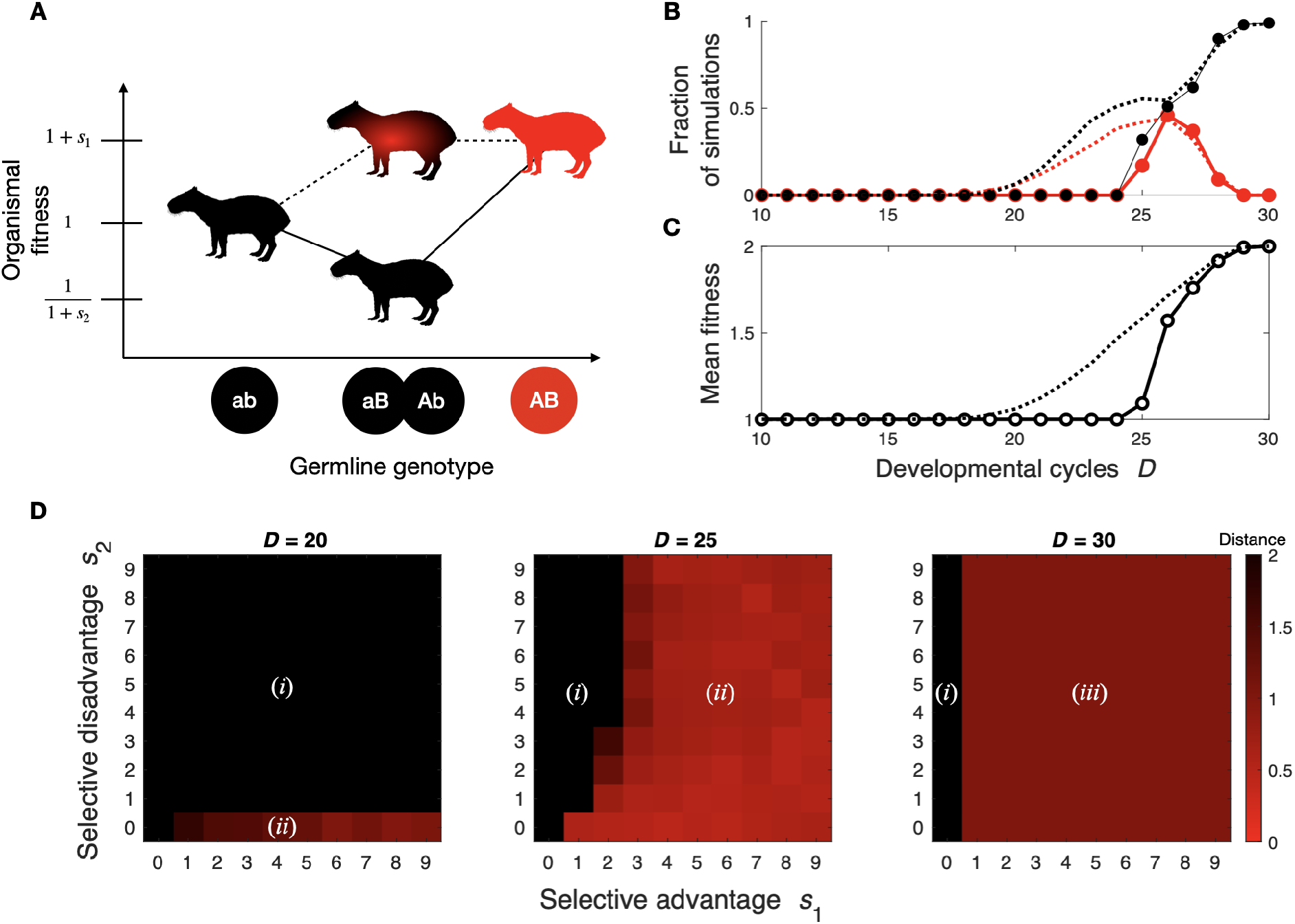
Somatic mutations can facilitate valley crossing. (A) We modified our baseline fitness landscape (Fig. 1A) to include a fitness valley at intermediate germline genotypes (i.e., at genotypes aB and Ab). (B) The fraction of simulations in which the population reached the peak genotype (red) or remained one mutation away from the peak genotype (black) after 5000 generations is shown in relation to *D*, for *s_2_* = 0.5. (C) Mean population fitness after 5000 generations for the same simulations as in (B). The dotted lines show the data from Fig. 3B and C, for reference. (D) Heatmaps of the mean distance to the peak after 5000 generations. Rows correspond to different values of the selective disadvantage *s_2_* of the genotypes aB and Ab, while columns correspond to different values of the selective advantage *s_1_* of genotype AB. The labels (*i*), (*ii*) and (*iii*) refer to the different evolutionary outcomes described in Fig. 3A. In all panels, *N* = 100000, *μ*_soma_= 5×10^−9^ and *μ*_germline_ = 1×10^−8^.

Using our baseline model with a fitness valley of *s*_2_ = 0.5, the same three qualitative evolutionary outcomes emerged as without the valley, but the parameter combinations yielding each outcome changed quantitatively (Fig. 5B). We further explored how different combinations of selective values for peaks *s*_1_ and valleys *s*_2_ affected the likelihood of converging on the peak under different somatic mutation supplies, by varying the number of developmental cycles (Fig. 5C). For a low somatic mutation supply (*D* = 20), populations never converged on the peak if there was a valley, in contrast to our baseline model without a valley (*s*_2_ = 0, Fig. 5D). For an intermediate somatic mutation supply (*D* = 25), populations converged on the peak so long as *s*_1_ was not too low. For a high somatic mutation supply (*D* = 30), populations remained one mutation away from the peak regardless of the depth of the valley, so long as *s*_1_ > 0. Thus, a high somatic mutation supply can completely mask the deleterious effects of the intermediate germline genotypes, such that evolving populations converge on genotypes that would be otherwise maladaptive in the absence of somatic mutations. In sum, this analysis shows that non-heritable somatic mutations can facilitate valley crossing, but whether or not they do depends on the depth of the valley and the somatic mutation supply.

## Discussion

Somatic mutations are abundant and sometimes increase organismal fitness [31–33]. Nonetheless, because of their non-heritability, they are typically neglected as a source of adaptation in evolutionary theory [6, 41]. Here we provide proof-of-principle that non-heritable somatic mutations can promote adaptation and help traverse fitness valleys. They do so by exposing adaptive genotypes to selection ahead of their emergence in the germline, thus channeling populations toward peaks in adaptive landscapes, a process we refer to as somatic genotypic exploration. Below, we discuss the biological plausibility of somatic genotypic exploration given the simplifications in our model and the restrictions it uncovered, as well as the implications of this process for adaptive evolution.

### Biological plausibility of somatic genotypic exploration

Our model makes assumptions that simplify several intricacies of organismal biology. We modelled organisms as haploid individuals with asexual reproduction, whose somas develop through consecutive symmetric and synchronized cell divisions. Complexifying the model could make somatic genotypic exploration more or less likely, depending on the circumstances. For example, if the organism was diploid, the chances of acquiring somatic peak genotypes would be doubled, because there would be two copies of each allele per cell, but if the peak allele is recessive, somatic mutations would be less effective in revealing adaptive phenotypes. Moreover, tissue architecture and growth dynamics can affect the fate of somatic mutants [69]. More realistic developmental models accounting for differentiation, asymmetrical divisions and stem cells, will likely affect how somatic genotypic exploration influences adaptation. For example, somatic mutations could have greater influence if they arise in stem cells that contribute substantially to the composition of a tissue, but if somatic mutations are beneficial only if they arise in cells with specialized functions, then acquiring the somatic mutations in specific developmental contexts where they are adaptive would be less likely.

Despite these simplifications, our model is suggestive of the biological conditions under which somatic genotypic exploration is expected to influence adaptive evolution in nature. For example, our model suggests that non-heritable somatic mutations can promote adaptation when the somatic mutation supply is high, which can occur by an increased number of cellular divisions in development and by an increased somatic mutation rate. One could object that an increased somatic mutation rate would cause deleterious mutations elsewhere in the genome, thus offsetting any beneficial effects of somatic mutations. However, mutation rates are highly heterogeneous across the genome, sometimes varying by several orders of magnitude even amongst neighboring loci [70, 71], which produces mutational hotspots that can be sources of evolutionary adaptations [72, 73]. Similarly, these hotspots may confer the elevated somatic mutation rates suggested by our model to promote adaptation via non-heritable somatic mutation, without increasing the mutation load elsewhere in the genome.

Our model also suggests that non-heritable somatic mutations can promote adaptation when only a small number of cells with the adaptive somatic mutation are required to confer a selective advantage to the organism. Cell signaling offers a promising example of a biological process where this may occur, because a small number of cells with a somatic peak genotype influencing signal emission could orchestrate the behavior of many more cells with alternative genotypes. For example, somatic mutations in organs with endocrinological functions can drastically alter individual physiology and development [74, 75], even when those mutations do not cause enhanced cell proliferation and are in normal non-tumoral tissue [76, 77]. In a developmental context, small disturbances from signals could trigger the formation of new patterns in embryos [78], disturbances which could come about from somatic mutations affecting paracrine signaling in few cells amongst a population of cells that do not have the somatic mutation. As an illustrative example, Maderspacher & Nüsslein-Vollhard [79] reported the rescue of a wild-type striped phenotype in mutant zebrafish incapable of producing stripes, enabled by a small number of wild type cells derived from grafting their progenitors into the mutant embryo.

Furthermore, according to our model, non-heritable somatic mutations are more likely to promote adaptation when they confer a selective advantage not only to the organism, but also to the somatic cells in their developmental context. In other words, somatic genotypic exploration is facilitated if somatic mutations that are beneficial to the organism also confer a competitive advantage to the cells in which they arise, allowing these cells to increase in frequency within the organism. Many morphological innovations in animal evolution resulted from changes in cell proliferation and the parameters controlling its onset and cessation [80, 81]. For example, differences in cell proliferation in the facial development of some species of phyllostomid bats help explain the different lengths of their snouts in relation to the shape of the flowers they feed on [82], and genes involved in cell proliferation show indications of positive selection in animals of relatively large size such as capybaras [83]. Conceivably, different degrees of cell proliferation revealing beneficial phenotypes like in these examples might have arisen first from prolific somatic variants, and selection acting on the potential to acquire those mutations eventually promoted their emergence in the germline.

### Evolutionary implications of somatic genotypic exploration

Somatic genotypic exploration can impact evolution in at least three ways. First, somatic genotypic exploration can make the vast supply of non-heritable genetic diversity adaptively relevant. As was pointed out by Frank [84], the cell lineage history of the development of a single human individual is tremendous, exceeding the lineage history of hominids. Given empirical rates of somatic mutations, a single body can thus harbor immense genetic diversity, which is only now starting to be explored in depth with sequencing technologies [23, 27, 31, 53, 85–88]. Such diversity cannot directly enter the germline, but by fueling somatic genotypic exploration it could still influence evolutionary trajectories, as our model shows.

Second, somatic genotypic exploration allows selection to act on the potential of genotypes to produce non-heritable adaptive phenotypes, eventually rendering those phenotypes heritable. This makes somatic genotypic exploration akin to the genetic assimilation of plastic phenotypes triggered by environmental conditions [89–92] or by the stochasticity or “noise” of cellular processes [54, 93–95]. Within this context, the so-called “look-ahead effect” [54] is particularly relevant; in this model, phenotypic mutations caused by transcription or translation errors create potentially adaptive protein variants, offering a mechanism for channeling populations towards adaptive genotypes, as in our model. However, because somatic genotypic exploration relies on the exploratory potential of somatic mutations that arise during development, it can act on substantially different phenotypes via substantially different biological processes. For instance, the effect of cell-lineage selection would be irrelevant in the absence of some degree of intra-organismal inheritance, which is provided by somatic mutations, but would be mostly absent in the transcriptional and translational errors enabling the look-ahead effect.

Third, somatic genotypic exploration can cause developmental bias. Developmental bias exists when certain phenotypes are produced more readily than others, thus influencing evolutionary trajectories and outcomes [96, 97]. They can arise either from developmental constraints impeding the emergence of certain phenotypes [98] or through developmental drive, which accounts for the increased likelihood of some phenotypes [99]. Some causes of developmental drive are high mutation rates in genomic regions affecting an evolving trait [72, 73], the genetic architecture of the trait [100, 101], and the number of genotypes mapping to a phenotype [102]. Somatic genotypic exploration is a form of developmental drive in the latter sense, because it causes genotypes to intermittently express beneficial phenotypes that they would not otherwise express in the absence of somatic mutation, thus altering the genotype-phenotype map.

Overall, our study offers a theoretical grounding for the further analysis of somatic mutations as a source of adaptation. Future empirical studies can help evaluate the plausibility of somatic genotypic exploration, through analyses of traits affected by genomic regions with high somatic mutation rates, phenotypes that can be altered by relatively few or clonally expanding mutant cells, and phenotypic innovations derived from changes in the proliferation or mortality of cells during development. For these analyses, we need to better understand the dynamics of somatic mutant cells within the organism and how somatic genetic diversity affects phenotypes beyond cancer and senescence. By studying somatic genotypic exploration as a potential adaptive mechanism, we can elucidate whether and how the immense genetic diversity of the soma directs evolutionary trajectories toward adaptation. If that proves to be the case, the soma and by extension the organism as a whole, is not only the instantiation of the founding genotype present in the zygote, but it is rather a cloud of pulsing genotypic potential that serves as a testing ground for evolutionary trajectories, so transcending a role as an evolutionarily absurd vehicle for genes.

## Methods

### Baseline model

We modelled a population of multicellular organisms with an impermeable germline-soma division navigating a fitness landscape. We used a two-locus two-allele model in which one of the four possible allele combinations represented the peak conferring a selective advantage *s*_organism_ over the other three genotypes (Fig. 1A).

We used Wright-Fisher simulations with a population of *N* haploid individuals, where we represented each individual by the mutational distance of its germline genotype to the peak genotype. We initialized monomorphic populations at the maximum distance from the peak (i.e., genotype ab, which is two mutations from the peak). We ran each simulation for 5000 generations, each of which consisted of a developmental phase and a reproductive phase (Fig. 1B).

In the developmental phase, we modelled the somatic growth of each individual in the population as a branching process with *D* developmental cycles consisting of synchronized rounds of cell divisions starting from a single cell, until the individual reached a reproductive somatic size 2^*D*^. The starting cell contained the germline genotype and at each cell division, somatic mutations occurred at rate *μ*_soma_ without altering the germline genotype. To implement this, at each round of cell division, we sampled the number of mutated cells from a binomial distribution B(*n, μ*_soma_), where *n* was the number of somatic cells with each genotype (ab, aB, Ab or AB). The genotypic composition of the soma at the end of development defined organismal fitness. In the case of our baseline model, having at least one somatic cell with the peak genotype provided the full selective advantage *s*_organism_, producing an organismal fitness of 1 + *s*_organism_; otherwise the fitness was 1.

In the reproductive phase, germline genotypes were selected with replacement from the population with a probability proportional to organismal fitness at the end of development. At this step, the germline genotypes of offspring were mutated at a rate *μ*_germline_ per locus. These selected and possibly mutated germline genotypes produced the population of the next generation.

As a control, we also considered a version of our model where somatic mutations did not affect fitness. To do so, we only included the reproductive phase in each generation. Organismal fitness was therefore defined exclusively by germline genotype.

The default parameters for our baseline model were *D* = 25, *N* = 100000, *s* = 1, *μ*_soma_ = 5×10^−9^ and *μ*_germline_ = 1×10^−8^. However, we also explored the parameter space by including ranges from *D* = 10 to *D* = 30, from *N* = 1000 to *N* = 200000, from *s* = 0 to *s* = 10, from *μ*_soma_ = 1×10^−10^ to *μ*_soma_ = 5×10^−3^ and from *μ*_germline_ = 1 × 10^−10^ to *μ*_germline_ = 5 × 10^−4^.

### Alterations to the baseline model

#### Fitness functions

We ran simulations in which organismal fitness was a function of *σ*_peak_, which is the fraction of the developed organism’s somatic cells with the peak genotype. To do so, we defined the fitness of each individual *i* as 1 + *s*_organism_*F*_i_(*σ*_peak_), where *F*_i_(*σ*_peak_) was a monotonic function of *σ*_peak_ that yielded values between 0 and 1. *F*_i_(*σ*_peak_) thus factored the selective advantage *s*_organism_ conferred to an individual, according to its fraction of somatic cells with the peak genotype. We defined *F*_i_(*σ*_peak_) as

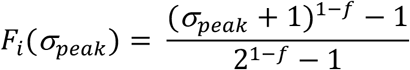

where *f* was a constant. We chose six values of *f* in order to study a range of fitness functions that required different numbers of somatic cells with the peak genotype to confer the full selective advantage *s*. These values were *f* = 0, 12, 107, 1055, 10538, and 10000 (Fig. 4A).

The set of baseline parameters we used in combination with these fitness functions was *D* = 25, *N* = 100000, a range of *s* from *s*_organism_ = 0 to *s*_organism_ = 10, *μ*_soma_ = 5×10^−9^, 1×10^−8^, or 5×10^−8^, and *μ*_germline_ = 1×10^−8^.

#### Cell fitness

We ran simulations in which somatic mutations to the AB genotype conferred a selective advantage not only to the organism, but also to the somatic cell. To do so, we added an extra stage to the developmental phase, in which somatic cells with AB genotypes had a proliferative advantage over somatic cells with other genotypes. Specifically, they divided at a rate *s*_cell_ times that of somatic cells with other genotypes. Somatic mutations occurred at the same rate *μ*_soma_ in these cell divisions as in other cell divisions. We used a value of *s*_cell_=2, approximating estimations for differential proliferative capacities among somatic variants [31, 58] and populations of cells in stages of development that are comparable to each other across different species [82].

#### Fitness valleys

We ran simulations in which the intermediate germline genotypes aB and Ab conferred a fitness disadvantage to the organism, relative to the genotypes ab and AB. Specifically, we modified our baseline model such that individuals with the ab germline genotype had a fitness of 1, AB germline genotype had a fitness of 1+ *s_1_*, and the intermediate germline genotypes had a basal fitness of 1/1+s_2_, which could be increased via somatic mutation to AB. In these simulations, we explored values for *s_1_* and *s_2_* ranging from 0 to 10 and values for *D* ranging from 10 to 30.

## Acknowledgements

We thank Steven A. Frank, David V. McLeod, and William S. Dewitt III, as well as members of the Computational Biology and Theoretical Biology groups in the Institute of Integrative Biology at ETH Zurich for critical feedback and discussions. This work was funded by the Swiss National Science Foundation (grant PP00P3_170604).

## Notes

### Competing Interest Statement

The authors have declared no competing interest.

